# Predicting treatment-free remission outcomes in Chronic Myeloid Leukemia patients using an integrated model of tumor-immune dynamics

**DOI:** 10.1101/2024.10.10.617526

**Authors:** Artur C. Fassoni, Agnes Yong, Richard E. Clark, Ingo Roeder, Ingmar Glauche

## Abstract

The interactions between tumor and the immune system are main factors in determining cancer treatment outcomes. In Chronic Myeloid Leukemia (CML), considerable evidence shows that the dynamics between residual leukemia and the patient’s immune system can result in either sustained disease control, leading to treatment-free remission (TFR), or disease recurrence. The question remains how to integrate mechanistic and data-driven models to support prediction of treatment outcomes. Starting from classical ecological modeling concepts, which allow to explicitly account for immune interactions at the cellular level, we incorporate time-course data on natural killer (NK) cell number, function, and their tumor-induced suppression into our general model of CML treatment. We identify relevant time scales governing treatment and immune response, enabling refined model calibration using tumor and NK cell time courses from different datasets. While the model successfully describes patient-specific response dynamics, critical parameters for predicting treatment outcome remain uncertain. However, by explicitly incorporating tumor load changes in response to TKI dose alterations, these parameters can be estimated and used to derive model predictions for treatment cessation. Further exploring dynamic changes in the number of functional immune cells, we suggest specific measurement strategies of immune effector cell populations to enhance prediction accuracy for CML recurrence following treatment cessation. The generalizability and flexibility of our approach represent a significant step towards quantitative, personalized medicine that integrates tumor-immune dynamics to guide clinical decisions and optimize dynamic cancer therapies.

## Introduction

Conventional chemotherapy has achieved substantial advancements in cancer treatment. However, the availability of targeted therapies opens a new spectrum of applications focusing on specific tumor features and their interactions with potentially supportive environments [1]. It is increasingly recognized that the patient’s immune system plays a crucial role in controlling tumor remission [2, 3]. While the immune system may not have initially suppressed tumor outgrowth, anti-tumor treatments can help reestablish this balance. Different immunotherapeutic approaches actively target this process to stimulate existing anti-tumor responses, for example, by enhancing recognition and cytotoxic capacities of T and natural killer (NK) cells [4]. Specifically, CAR-T cell-based therapies modify patient derived T cells with artificially designed receptors to recognize and actively destroy tumor tissues [5, 6].

Many immunotherapeutic approaches, however, suffer from high inter-patient variability and severe side effects [7, 8]. The poor predictability of therapy response often stems from an incomplete understanding of the complex interactions involving the immune system and how it can be effectively guided or activated. Moreover, this process is not unidirectional: tumor cells can also strongly alter or even disable immunological control mechanisms [9].

In this context, we recently suggested that monitoring the changes in disease dynamics in general may provide critical insights into treatment outcomes [10]. Furthermore, there is increasing evidence that changes in the number and function of immune cells over time can provide important information about an individual’s immune response [11–13]. Within the paradigm of personalized medicine, it is therefore conceivable that patient-specific treatments schedules, which adapt to the dynamic response of tumors, will also incorporate the evolution of critical immune components to guide further treatment decisions [14]. The future potential of such applications extends beyond tumor targeting and control to include tumor prevention [15]. Therefore, it is vital to develop a quantitative understanding of the various ways in which immunologically relevant cell populations interact with tumor tissue and its particular microenvironment.

Chronic myeloid leukemia (CML) developed into a primary example to illustrate the enormous potential of targeted therapies, altering CML into a “controllable disease”. As a result of this success, most patients with CML have an almost normal life expectancy [16]. Although continuous treatment with tyrosine kinase inhibitors (TKI) is highly efficient, life-long therapy is associated with side effects, a lower quality of life, and high economic costs. Therefore, treatment cessation for optimally responding patients, leads to a characteristic outcome: About half of the patients remain in treatment free remission (TFR) while the other half presents with CML recurrence usually within 12 months after therapy stop [17–19]. It has been suggested that immunological mechanisms are a central determinant to account for the particular patient outcome [19–23]. This is especially prominent as some patients show low but non-expanding leukemia levels, which can hardly be explained without an immunological control mechanism [24]. While a number of studies have suggested that the number and function of various immune cell types at the time of treatment cessation may serve as predictive markers for TFR, there is currently no consensus on this question.

Mathematical and computational models of CML are an essential tool to obtain a quantitative understanding of the mutual interaction dynamics between different relevant cell populations [25–39]. Although first CML models with an explicate consideration of leukemia-immune interactions emerged rather early [35, 36], it was only until the TKI cessations was considered as a relevant treatment alternative for optimally responding CML patient that this particular aspect received more attention, especially to provide a conceptual understanding about why some patients remain in treatment-free remission, while other patients are relapsing [37–39].

We have formally demonstrated that the different potential states after treatment cessation - i.e., recurrence, disease control, and cure - impose specific requirements for the interaction between leukemia and responsive immune cell populations [37]. Additionally, we have shown that altering treatment schedules, such as dose reductions, can provide informative readouts to “probe” the immunological configuration and identify different risk groups [38–40]. However, these phenomenological estimates do not necessarily reflect the mechanisms at the cellular level underlying immune-leukemia interactions. Considering increasing efforts to identify and prospectively measure relevant immune populations, it is crucial to understand how a decreasing leukemia load translates into an activated immune response. Knowledge of these mechanisms will enable the estimation of patient-specific parameters and potentially translate them into optimized predictions for the patient’s response to treatment cessation.

In this work, we integrate mechanistic and data-driven approaches to describe tumor-immune interactions and predict TKI treatment cessation responses in CML patients. Revisiting classical ecological models of predator-prey interactions, we explicitly account for cellular-level mechanisms, such as searching, targeting, and recharging of immune effector cells, as well as the tumor-induced suppression of immune activity. This modeling approach provides a mechanistic basis for the concept of the “optimal immune window” in tumor control [36, 37]. By incorporating NK cell dynamics from published time-course data [13], we quantify tumor-induced suppression of immune recruitment and function, identifying characteristic time scales in model dynamics that delineate different dominating effects. Fitting the model to time courses of 75 patients from the DESTINY trial [39, 41], we show that measuring the dynamic response in tumor load after dose reduction is essential to correctly estimate the leukemia-immune parameters necessary for predicting the effect of treatment cessation. In a complementary manner, we also show that measuring the dynamic change in the number of functional immune effector cells provides predictive markers to anticipate treatment outcomes. Our generalizable model provides a fundamental proof-of-concept step towards personalized-medicine approaches that integrate tumor and immune cell dynamics to guide treatment decisions.

## Results

Our current approach to dynamically model tumor-immune interactions in CML patients builds on earlier models of hematopoietic stem cell organization and leukemia treatment [26, 30, 37, 39, 42]. Those models assume two states for (tumor) stem cells, namely a proliferative, activated state, and a non-proliferative, quiescent state. Because there is long-standing evidence that quiescence of (tumor) stem cell is reversible [43] and mediated by a specific microenvironmental context, the so-called stem cell niche [44, 45], we denote the quiescent tumor (stem) cells also as niche-bound. Within the modeling framework, the population of tumor cells interacts with a population of immune effector cells (Fig. 1A, see Methods). The current analysis focuses specifically on three mechanistic aspects tumor-immune interactions: the antitumor activity of immune cells (represented by *f*(*T*)), and the suppression of immune cell number and function (described by *g*(*T*) and *h*(*T*), respectively).

**Figure 1:**
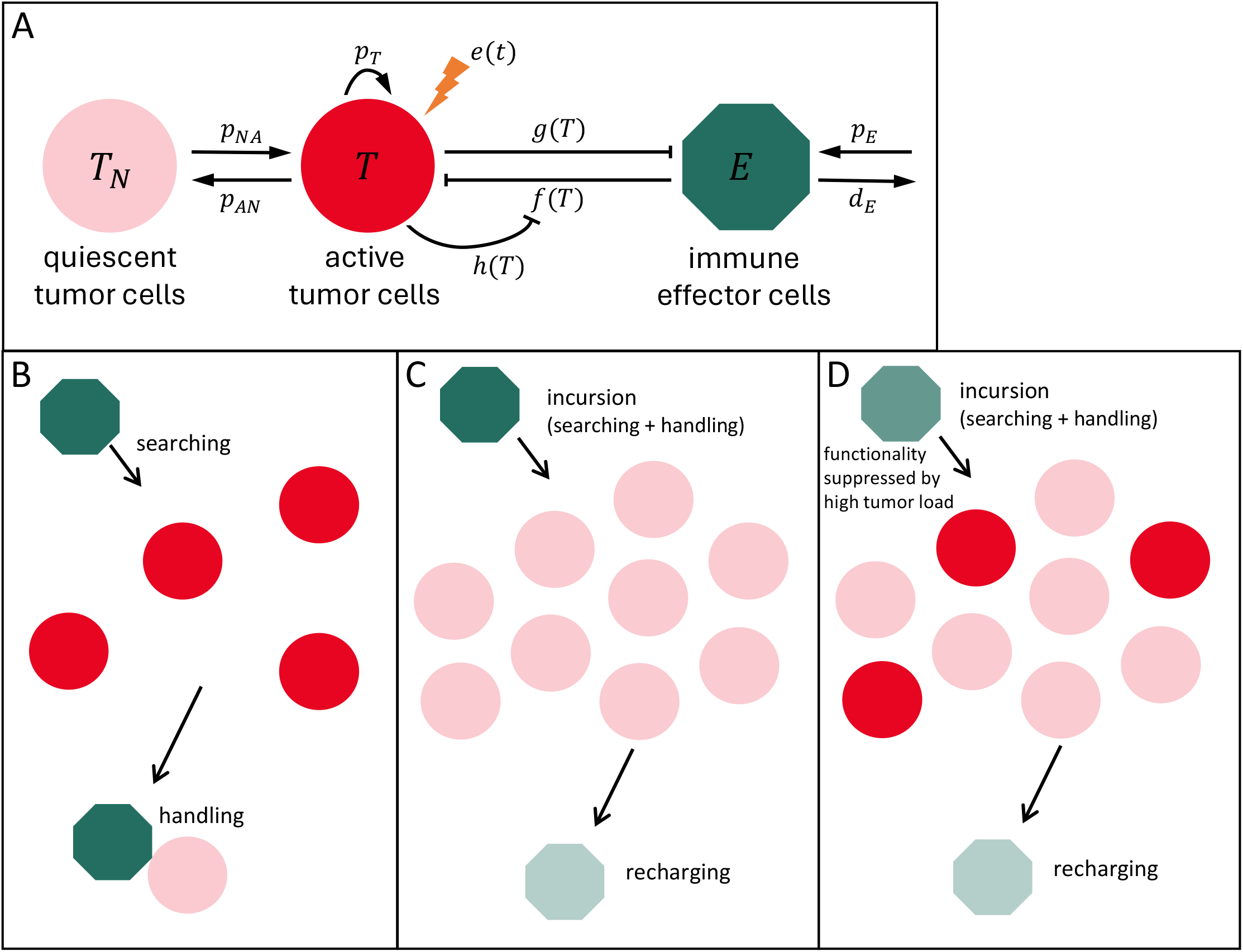
Model for tumor-immune interactions. A) The model assumes two states for tumor cells, namely active *T* and quiescent, niche-bound tumor cells *T*_*N*_, and a compartment of immune effector cells *E*. Tumor cells in each state transition to the other state with rates *p*_*NA*_ and *p*_*AN*_. Only the active tumor cells proliferate with rate *p*_*T*_, are targeted by therapy with rate *e*(*t*), and interact with immune effector cells. These are produced at a constant rate *p*_*E*_, die with rate *d*_*E*_ and kill tumor cells with immune effector activity *f* (*T*). Tumor cells suppress the recruitment and function of immune cells, as described by *g*(*T*) and *h*(*T*). B,C,D) Modeling immune cell activity. A functional response describing immune cell-mediated tumor killing is derived assuming mechanisms of searching, handling, and recharging. B) Similar to predator-prey approaches, immune cells (green) act as predators that perform incursions to hunt prey (tumor cells, red), consisting of a search phase (with duration inversely proportional to prey density) and a handling phase in which the attached cell spends a fixed amount of time to destroy the prey (killed tumor cells in light red). C) We further assume a recharging phase in which the now exhausted immune cell (light green) spends a fixed amount of time regaining the effector phenotype. D) Consistent with recent findings, we assume that functionality of immune effector cells is suppressed by a high density of tumor cells, leading to reduced efficiency in eliminating tumor cells.

### The immune response can be explained by a superposition of a searching, handling and recharging process and its inhibition by tumor load

To model the antitumor immune activity *f*(*T*), we assume that the targeting of tumor cells is determined by a process at the individual immune cell level, where immune cells perform the serial killing of tumor cells during an “incursion” consisting of three phases (Fig. 1B,C) [46-52]: i) *searching* for tumor cells, spending a searching time inversely proportional to the tumor load, ii) *handhing* of tumor cells, with a fixed time interval in which the immune cells are attached to a tumor cell, and iii) a *recharge* phase, where immune cells recover their original functional status after having completed the searching and handling sequence a certain number of times. This recharging of immune competence is required before the incursion can begin again. Incorporating these aspects and revisiting classical predator-prey interactions, we show (see Methods) that the antitumor activity resulting from these processes is described by a Holling type-2 functional response

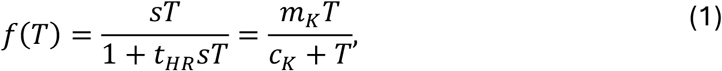

with saturation level *m*_*K*_ = 1/*t*_*H*_ and half-saturation constant *c*_*K*_ = 1/(*t*_*H*_*s*) related to the mechanistic parameters *t*_*H*_ (handling and recharging times) and *s* (search speed).

This description conveys the normal antitumor activity of immune effector cells, but it is known that their function is inhibited in CML patients at diagnosis and is restored when treatment decreases the tumor load [13, 20, 53]. To reflect a reduced functionality of immune cells as a function of tumor load, we multiply the antitumor activity by *h* ∈ [0,1], implying that only a fraction of all encounters is effective (Fig. 1D). A model for *h*(*T*) is

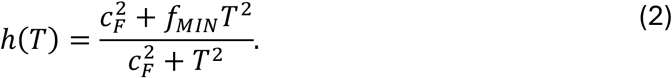

This starts at *h*(0) = 1, reflecting a full immune functionality in the absence of tumor cells, decreasing with increasing tumor load *T* towards reaching a minimal level *h*(∞) = *f*_*MIN*_ ≥ 0, indicating a suppressed immune function for high tumor load. Using the percentage of CD107a^+^ NK cell as a surrogate for NK cell function at different tumor loads shows that (2) provides an excellent fit and model for immune function (Fig. 2A).

**Figure 2:**
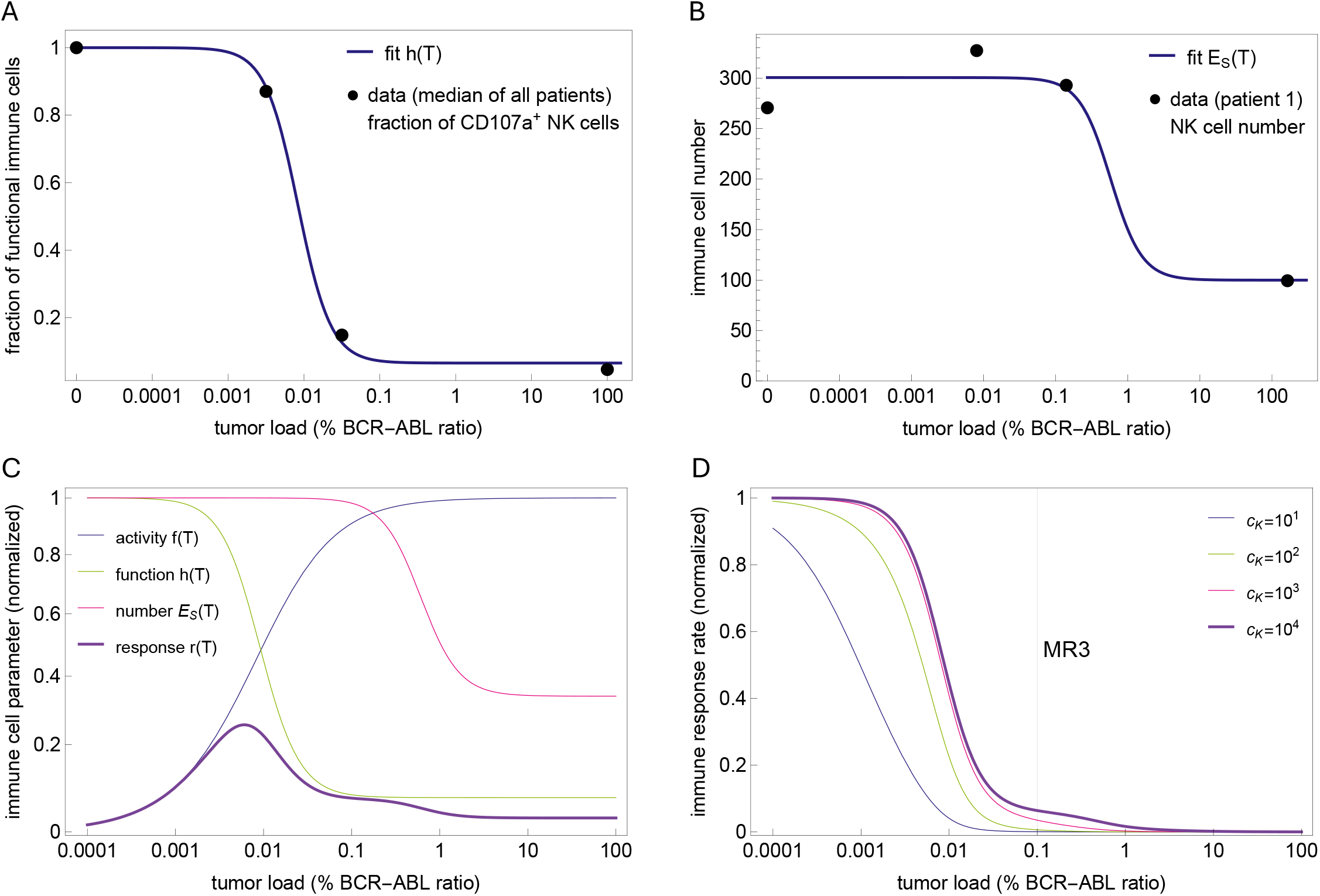
Estimating immune parameters from NK cell function and number. A) Using the % of CD107a+ NK cells as a surrogate for NK cell functionality, the model assumes that the immune effector function is suppressed at high tumor load, as seen in the data and described by the fitted function *h*(*T*) (equation (2)). B) NK cell data shows that immune cell numbers are decreased at diagnosis and recover as treatment induces tumor reduction. The model assumes that this suppression is described by *g*(*T*) (equation (3)). Since the dynamics of immune cells occurs on a fast time scale (Fig. S1A), it is possible to use a quasi-steady approximation to directly relate time courses of immune cell numbers and tumor loads with equation (4) and fit its parameters. The fit for a particular patient is shown here; the fits for the other 7 patients are shown in Fig. S1B-H. C) Plots of normalized immune cell functional responses *f* (*T*)/*f* (0), *h*(*T*)/*h*(0), *E*_*S*_(*T*)/*E*_*S*_(0) and *r*(*T*)/*r*(0). The effective immune response *r*(*T*) (equation (5)) represents the total number of tumor cells removed per unit time and depends on the actual tumor load. As a product of increasing and decreasing functions (*r*(*T*) = *f* (*T*)*h*(*T*)*E*_*S*_(*T*)), it has a bell shape describing an optimal immune window where the antitumor immunological effect is maximal. D) Plot of the normalized immune response rate, *k*(*T*)/*k*(0) (equation (6)). This is a dynamic rate that is modulated by the tumor load, changing from nearly zero at diagnosis at high tumor loads to its restored, original level *k*(0) at low tumor loads. There is a threshold for the tumor load that defines the sudden transition from zero to the plateau level. Although this threshold depends on the parameter *c*_*K*_, testing different reasonable values shows that this threshold is never above MR3.

We conclude that the immune-mediated killing of tumor cells results from a superposition of the density-dependent searching-handling-recharging process with a tumor-dependent inhibition of immune cell functionality.

### Restauration of immune cell numbers can consistently be explained as a response to therapy-driven reduction in tumor load

In addition to their functional impact, tumor cells have been shown to suppress the number of immune cells [12, 13, 21]. This effect is reversible, as demonstrated by a recovering of immune cell number during therapy [13]. Most likely, this is not a direct effect of the therapy, but results from the reduction in tumor load. In our model, the tumor-dependent modulation of immune cell recruitment/production is described by

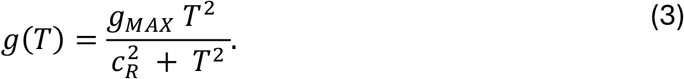

This formulation is inspired by our previous comparison of different models for immune recruitment [37]. It assumes that the suppression of immune cell number is small for few tumor cells, increases as the number of tumor cells increases, and saturates to a maximum *g*_*MAX*_, reaching half of value when *T* = *c*_*R*_.

To assess the compatibility of this function with NK cell data, we observe that regulation of immune recruitment occurs on a fast time scale (Fig. S1A). Thus, in a short time interval, the number of immune cells reaches the quasi-steady state *dE*/*dt* = 0, called slow manifold, given by equation (10) in Methods. Substituting *g*(*T*) in (10) we obtain a direct relationship expressing immune effector cells *E* as a function of the tumor load *T*,

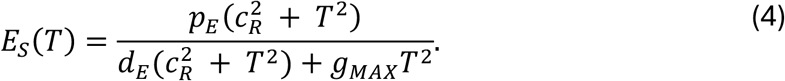

The published dataset (from [13]) we used at this point, includes eight time courses with points (*T*_*i*_, *E*_*i*_) consisting of NK cell counts *E*_*i*_ and tumor load *T*_*i*_ at different time points. For each patient, we found the parameters *p*_*E*_, *c*_*R*_, *g*_*MAX*_ that fit (4) to the data (*T*_*i*_, *E*_*i*_) (Figs. 2B, S1B-H), showing that (3) is an adequate model and further illustrating that the therapy induced reduction of tumor cells leads to the restoration of normal immune cell numbers.

We conclude that monitoring of immune cell numbers and using the model timescale separation allows to estimate the shape and parameters for the functional response *g*(*T*) describing the tumor-mediated suppression of immune recruitment.

### The model reveals an optimal window and a scale separation for the immune response

Our modeling approach allows estimating the functional shape of tumor-immune interactions, but also provides a mechanistic explanation for the so-called “immune window” [3C, 38]. Indeed, the reduced model (equation (11)) shows that, along the slow manifold, the antitumor immune effect as a function of the tumor load is described by

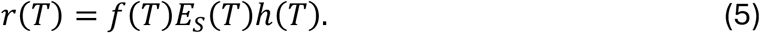

In other words, the effective antitumor immune response *r*(*T*) is the product of immune activity (*f*, increasing with *T*), the total number of immune cells (*E*_*S*_(*T*), decreasing with *T*) and the fraction of functional immune cells (*h*, decreasing with *T*). This convolution of increasing and decreasing functions leads to a bell shape for *r*(*T*) and to the emergence of an “optimal immune window”, in which the antitumor immunological effect is maximal for intermediate ranges of the tumor load (Fig. 2C). Intuitively, this results from the rare encounters with tumor cells at low tumor loads when the search process dominates (small *f*), while the immune function and recruitment are suppressed at high tumor load (small *h* and *E*_*S*_). Although such an “immune window” description was used before [3C, 38], it was assumed *a priori*, whereas here it emerges from explicit modeling of the relevant immunological mechanisms and is further supported by NK cell data.

While the antitumor effect *r*(*T*) describes the number of tumor cells removed by time, the *eflective immune response rate* describes the *per capita* rate of this removal at the overall population level, and is given by

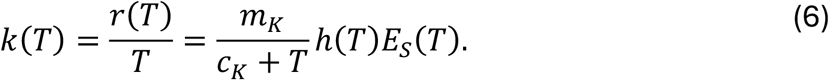

Similar to the constant TKI effect and the tumor growth rate, the immune response rate is given in units of 1/*time*, being, however, a dynamic kill rate changing in different scales in response to the tumor load (Figs. 2D, 3AB). Indeed, it appears essentially as a step function with two scales: at a high tumor load, it is negligible compared to the proliferation and TKI kill rates, while at tumor load below a certain threshold, it reaches a plateau level close to its maximum value *k*(0) = *p*_*E*_*m*_*K*_/(*d*_*E*_*c*_*K*_). Importantly, while the plateau level depends on the ratio *m*_*K*_/*c*_*K*_ between the saturation level *m*_*K*_ and the half-saturation constant *c*_*K*_ for immune cell activity, the threshold at which *k*(*T*) is restored does not depend on *m*_*K*_. As shown in the Methods, the NK cell time courses allow to estimate all except these two parameters of immune effector cells. However, testing different reasonable values for *c*_*K*_ shows that this threshold is never above MR3 (Fig. 2D), because at this tumor load the immune cell number and function are already suppressed to their minimum. This allows us to conclude that, independent of the two unknown immune cell parameters, the immune response rate is negligible for tumor loads at the scale above MR3 due to the suppression of immune cell number and function.

In summary, our modeling approach explains the immune window as an interplay of different immunological mechanisms and reveals a scale separation for the immune response rate, showing that it is only effective after the tumor load is reduced below MR3.

### The model describes patient time courses, but TFR prediction requires dynamic response data

The quantitative model reveals a scale separation for tumor-immune interactions which allows us to group model parameters into distinct modules and estimate them in a stepwise fashion. In brief, the step-function-like behavior of the effective immune response rate enables the estimation of all but two model parameters by first fitting a simplified model without the immune system and then fitting the remaining two with the full model (see Methods).

With this approach, while the other model parameters are constrained by the tumor time course under treatment, the degrees of freedom associated with response to treatment cessation are captured by the remaining parameters *p*_*T*_, *m*_*K*_, related to tumor growth and immune cell activity. Estimating these parameters using the full model and time courses under full dose only, leads to parameter combinations that provide equally good fits but predict different outcomes after therapy stop (Figs. 3CD, S2). Therefore, while these parameters are critical for predicting TFR, they cannot be inferred from the available time courses of immune cells and tumor load under standard therapy. In other words, these data do not contain the patient-specific signature quantifying how strong the immune system is at its restored level, or how fast tumor cells are growing at low densities.

**Figure 3:**
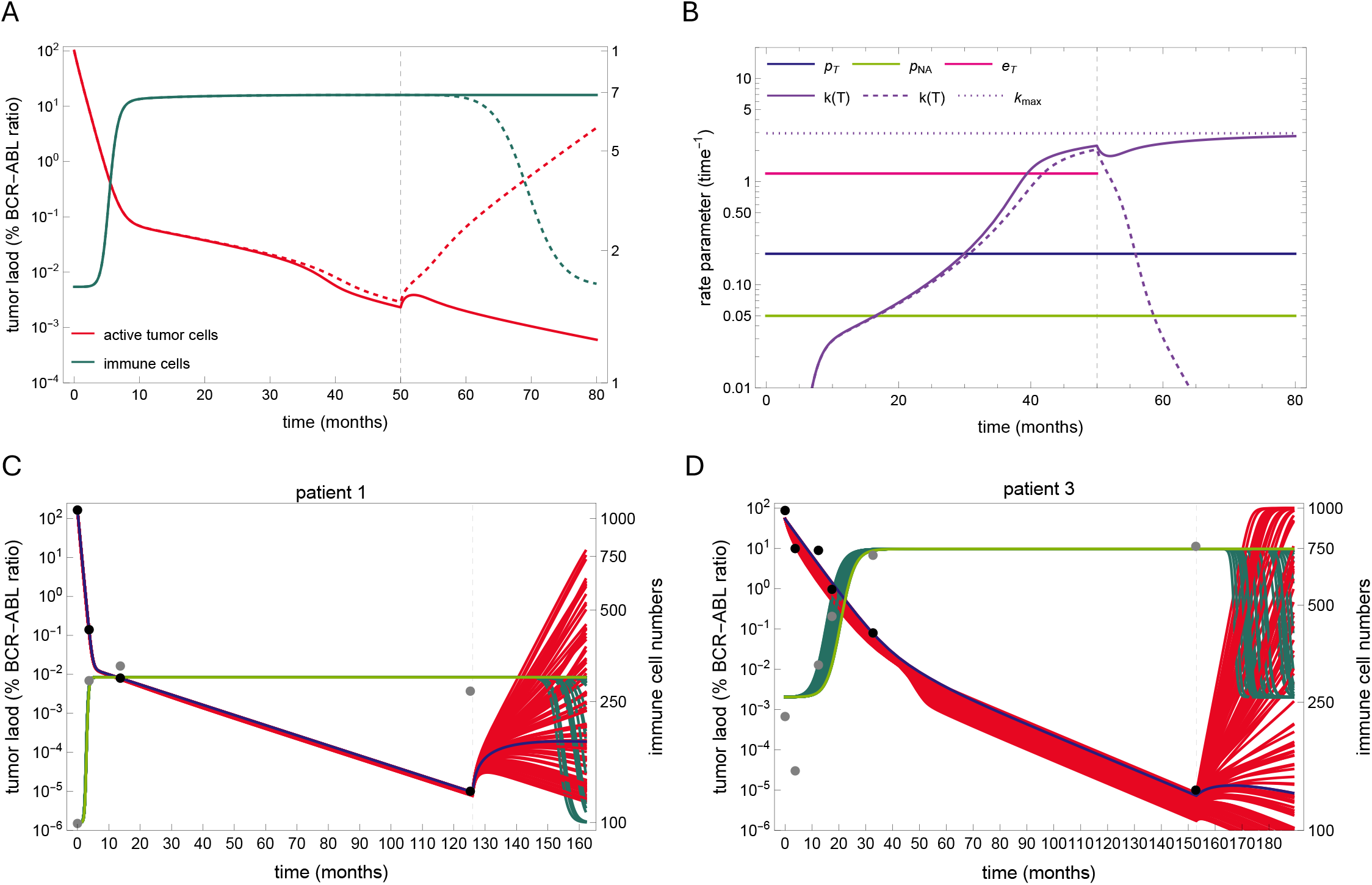
Model dynamic behavior. A) Two model simulations with identical representative parameter values except for the maximum immune activity *m*_*K*_ show different outcomes without any difference in the time course under standard therapy. After treatment cessation (dashed vertical line), the residual tumor cells may exhibit different responses, ranging from transient growth controlled by immune cells (red solid line) to relapse (red dashed line). B) Comparison of constant rate parameters (*p*_*T*_, tumor growth rate; *p*_*NA*_, quiescent tumor cell activation rate; *e*_*T*_, treatment-induced tumor kill rate) and the dynamic immune response rate *k*(*T*) (purple curves). This dynamic rate starts from zero at diagnosis and is slowly restored until it reaches a level close to its maximum *k*_max_ = *k*(0) (dotted line). After treatment cessation, in the case of remission (purple solid line), the transient growth of tumor cells initially decreases the immune response rate, but it is restored after the control of the residual tumor cells, while in the case of relapse (purple dashed line), the tumor growth suppresses the immune response. C,D) Model fits for two representative patients. Model simulations of the 100 best fits (red for tumor cells, green for immune cells) to the time courses of tumor load (black dots) and NK cells (gray dots). Although the fits are indistinguishable under full-dose treatment, they predict different outcomes after therapy cessation. The best fit is shown in purple for tumor cells and light green for immune cells. See Figure S3 for the fits of the other six patients.

A way to overcome this limitation is to challenge the immune system against the residual tumor cells in order estimate its parameters. Dose alterations are a possible way for a clinical assessment of this relevant information. To illustrate this idea, we applied the model to a subset of 75 patients from the DESTINY trial [41], for which complete time courses before and during dose reduction are available [39]. Each patient underwent full-dose TKI therapy, followed by one year on a 50% reduced dose, before complete treatment cessation. Thus, we repeatedly applied the fitting procedure to these 75 patients, fitting *p*_*T*_ and *m*_*K*_ using data with increasing time intervals: i) full dose only, ii) full and reduced dose, iii) full and reduced dose and six months after treatment cessation. To account for the inherent non-uniqueness of these parameters, we selected the 100 best fits from 10,000 pairs (*p*_*Y*_, *m*_*K*_) for each setting. We then used these fits to predict the effect of treatment reduction and cessation on sustained remission, defining the patient-specific probability of molecular relapse at 12, 24 and 36 months as the fraction of non-remission fits that predict an increase in tumor load above MR3.

While the model predictions do not agree with the actual data when using the data under full dose only (Fig. 4A), including the data under reduced dose increases the prediction quality (Fig. 4B). Further including the additional response six months after treatment cessation leads to very good predictions (Fig. 4C). This confirms that the kinetics of tumor load after dose change contains information on the balance between tumor growth and immune response that was not available before, in line with previous suggestions [10].

**Figure 4:**
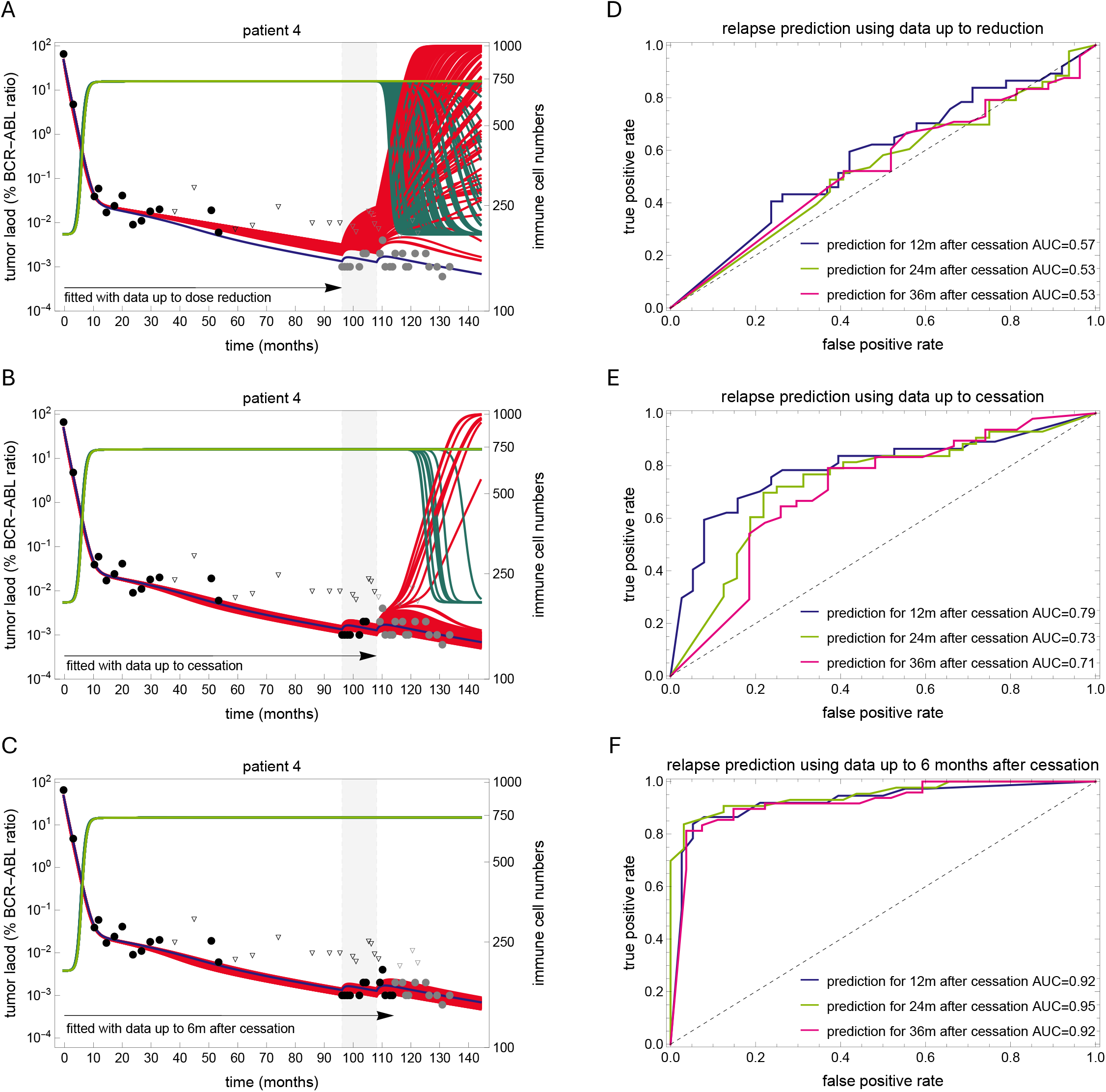
Model predictions with different data selections. A,B,C) best fits of the model (red for tumor cells, green for immune cells) obtained with different data selections for two representative patients. In each panel, the reduced dose period is shown with a light gray background, the data used for fitting are shown as black dots (triangles for undetectable values), while the data not used are shown in gray. The best fit using the full time course is shown in purple for tumor cells and light green for immune cells, and is used as a reference to compare the predictive ability of each model. The first selection (A) uses only the full-dose time course; the second (B) uses the full-dose and reduced-dose time courses, while the third (C) includes an additional six-month post-treatment period. D,E,F) Receiver operating characteristic (ROC) curves for each data selection setting. See also Figure S3.

We conclude that our model consistently describes both the tumor and immune cell time courses of CML patients, but that the essential information for predicting TFR is not encoded in the time courses under full-dose therapy and requires dynamic responses under different dose regimens.

### Dynamical changes in the number of functional immune cells predict relapse

We showed that the derivation of model predictions for TFR requires measuring tumor responses to dose alterations. Knowing the model mechanisms, we investigated whether monitoring immune cell number and function could additionally support TFR predictions. In this respect, recent efforts to establish immunological markers for TFR in CML suggest that NK cell subpopulations have predictive value [12, 13, 20–23].

We investigated the potential of the measuring the number of functional immune cells, which in the model is defined as the product of immune cell number (*E*) and function (*h*(*T*)). In the clinical context, this quantity can be calculated from NK cell time courses (like those in Fig. 2) by multiplying the absolute number of NK cells by the percentage of CD107a+ NK cells. We derive model predictions for the number of functional immune cells at different time points as follows. First, we restricted our analysis to the 100 best fits obtained with the complete time course of tumor load of each patient, as these estimates better capture the patient-specific parameters. The fits were split in two response groups, according to a predicted molecular relapse or persistent remission 12 months after therapy cessation. Then, we calculated the mean value of the number of functional immune cells across all fits in each group for each patient.

We found that the remission group corresponds to slightly higher values of functional immune cells at different times before and at treatment cessation, whereas stronger differences between the two response groups appear after cessation (Fig. S4). This suggests that a higher number of functional immune cells before treatment cessation associates with better response. However, we acknowledge that it might be difficult to anticipate the predictive information from single time points only.

Since the response of immune cells occurs on a faster time scale, we wondered whether the relative change in the abundance of functional immune cells after dose reduction correlates with treatment response, rather than the absolute values at a given time point (Fig. 5). We found that the loss of functional immune cells from dose reduction to 6 and 12 months thereafter strongly correlates with relapse, with losses larger than 20% highly associating with relapse (Fig. 6). On the other hand, smaller losses (reflecting a stable number of functional immune cells) are mostly associated with remission. However, this is not always the case, since some patients with a high proportion of relapse predictions do not show any loss during the reduced dose period. We found that most of these patients who relapsed later did present a loss of functional immune cells just 1 to 3 months after treatment cessation, indicating cases where the immune cell functionality took longer to deteriorate in response to the effects of dose reduction (compare panels in Fig. 6). In other words, as the tumor-immune interactions respond to dose alterations, a loss in the number of functional immune cells captures the essential information for reliable predictions and associates with relapse occurrence.

**Figure 5:**
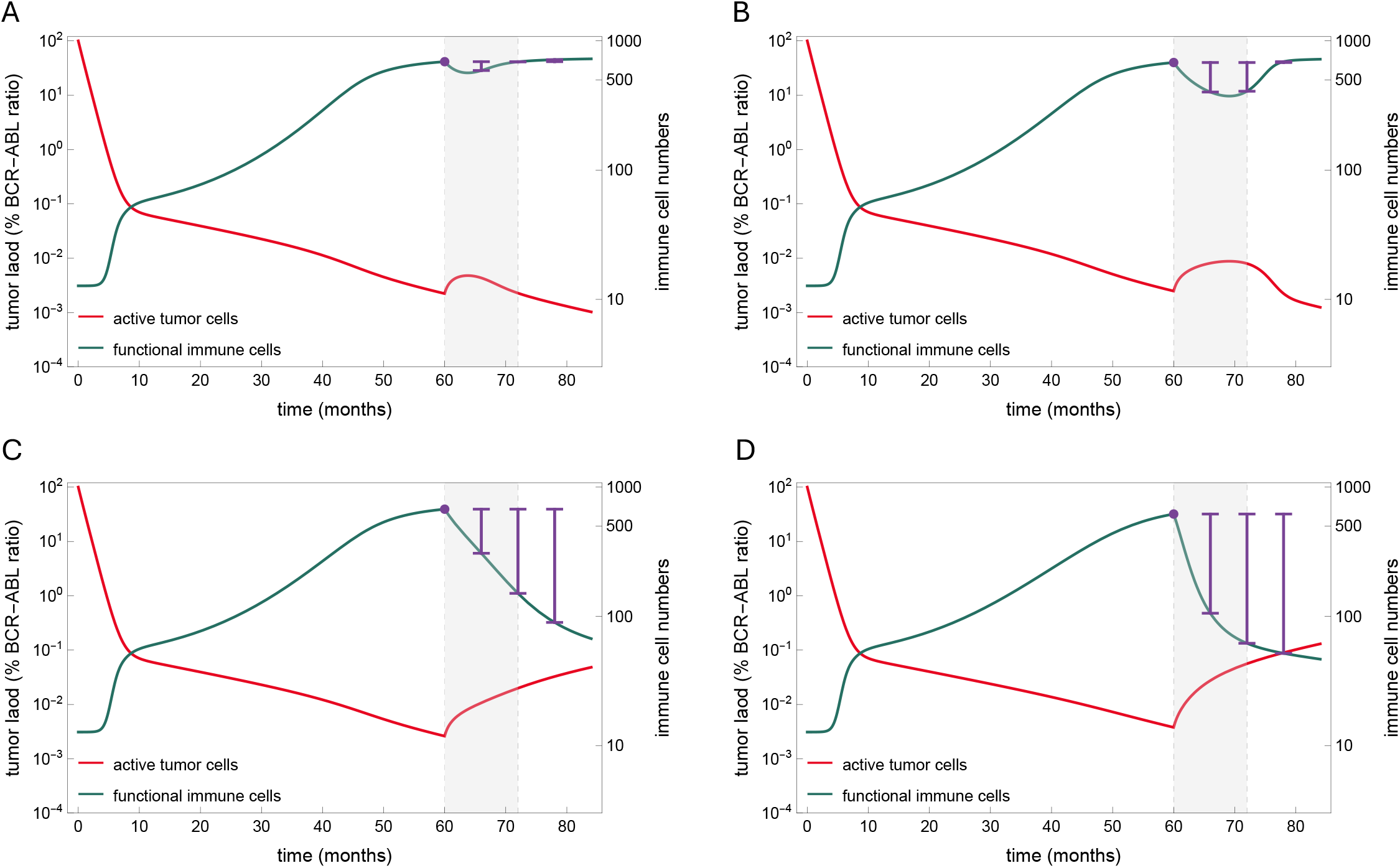
Dynamic changes in the number of functional immune cells after dose reduction to predict relapse. Model simulations showing similar tumor dynamics (red) during full dose treatment and different outcomes after dose reduction (month 60) and treatment cessation (month 72); the reduced dose period is shown with a light gray background. The number of functional immune cells, given as the product of immune cells *E*(*t*) and their functionality *h*(*T* (*t*)) is plotted in green. The purple lines illustrate how assessing the loss in the number of functional immune cells after dose reduction (reference value indicated by the purple dot) can predict the outcome of treatment cessation. Comparing the loss six months after reduction, a low value (A) is associated with remission, a high value is associated with relapse (D), while moderate values may be associated with both outcomes (B and C). However, evaluating the losses at later time points allows a better distinction between two cases, with a decreased value for remission (B) and an increased value for relapse (C).

**Figure 6:**
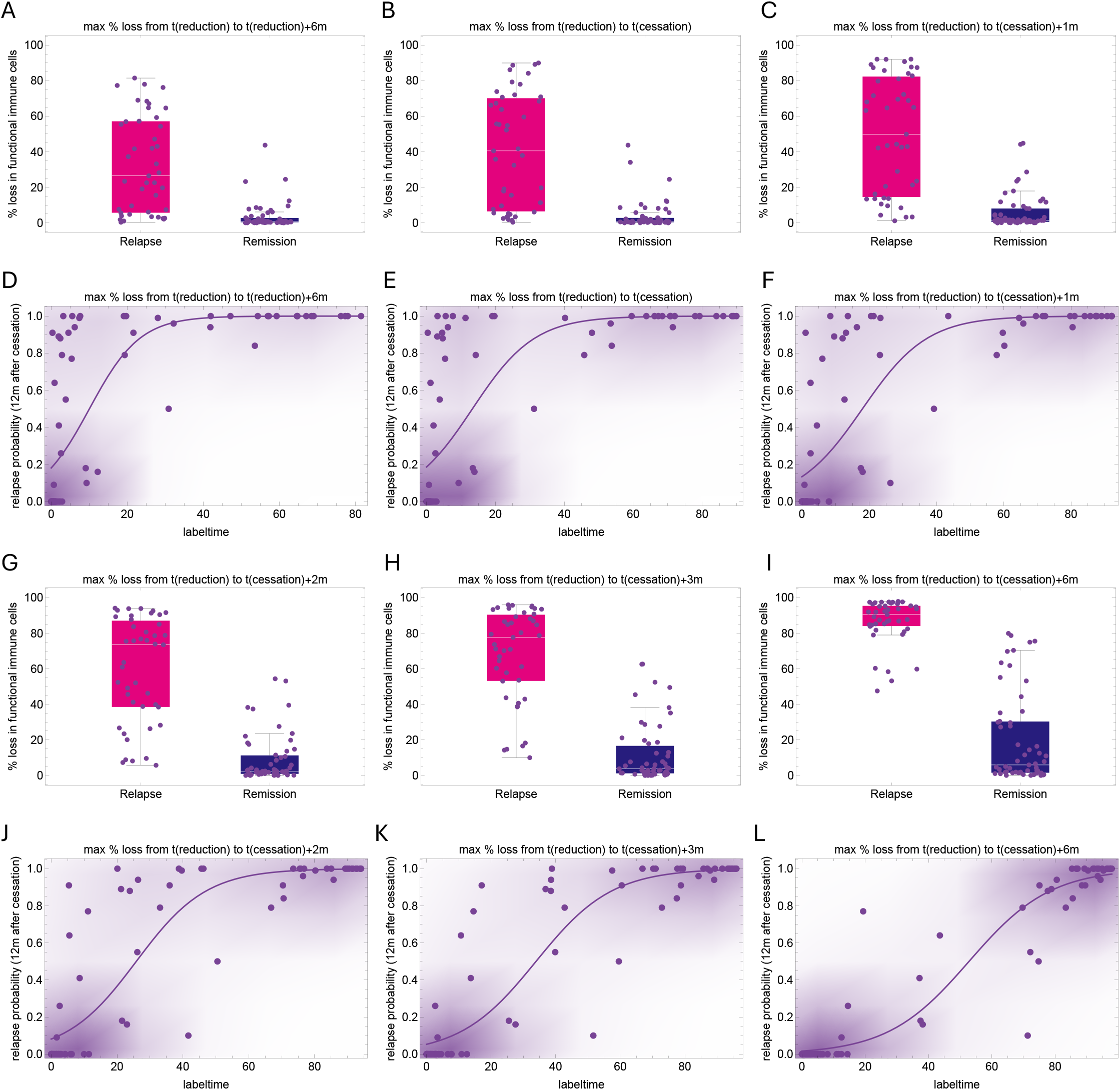
The dynamic change in the number of functional immune cells as a marker of relapse. A-C,G-I) Plots showing the distribution of model estimates (mean values across all best fits) for the patient-specific maximum % loss in the number of functional immune cells from the time of reduction to different later time points. D-F, J-L) Scatter plots showing the model estimates of the patient-specific probability of relapse and the maximum relative loss in the number of functional immune cells (mean of all best fits) from the time of reduction to to different later time points. A relative loss in the number of functional immune cells equal to or higher than 20% (decreased functionality) six months after reduction (A,D) is highly indicative of relapse, while losses close to 0 (stable functionality) may indicate a low probability of relapse or a later relapse. Waiting until treatment is discontinued (B,E) results in a stronger signal on the loss of functional immune cells, which is more pronounced at one (C,F) or more months later (G-L), allowing to distinguish those patients with initially stable immune functionality that was subsequently lost.

These results provide strong theoretical evidence that individual TFR success can be predicted by monitoring dynamic changes in the number of functional immune cells in response to dose alterations.

## Discussion

By extending our previous mathematical models of CML treatment, we here introduced a novel mechanistic model framework to describe tumor-immune interactions in CML patients by integrating different functional responses and cellular interactions. By explicitly incorporating NK cell dynamics, we suggest that immune-mediated reduction of tumor cells results from the combined effects of a density-dependent searching-handling-recharging process and tumor-induced inhibition of immune cell functionality and recruitment. As a main result, our model explains the concept of an optimal immune window as the consequence of these multiple interacting immunological mechanisms. While this concept has previously emerged as a phenomenological model, here we provide a mechanism-based foundation for it.

Our modeling results revealed a scale separation for the tumor-immune interactions, allowing to disentangle the underlying processes and showing that the immune response becomes effective only when the tumor load is sufficiently reduced, and thus can be analyzed separately from the initial response. Combining this knowledge with the fact that the three datasets employed in our approach - tumor load (BCR-ABL1 ratios), immune cell numbers (CD56dim NK cell counts), and immune cell function (CD107a+ expression) - are distinct in nature, we were able to link each dataset to a distinct module with a subset of model parameters and apply a tailored estimation procedure for each.

Consistent with our earlier results [37–39], we demonstrated that BCR-ABL1 dynamics under full-dose TKI treatment alone do not provide enough information to uniquely estimate critical immune effector cell parameters and to predict TFR success. However, we showed that including dynamical response data, such as BCR-ABL1 under reduced dose or following TKI discontinuation, offers a strategy to capture the additional information needed to identify these missing parameters [10]. Even more, we showed that further incorporating the dynamical response of NK cells counts to dose alterations, we can significantly improve the model’s predictive power regarding the success of treatment cessation. Therefore, we suggest measurements of immune effector cell properties that could serve as predictive markers for relapse after treatment cessation. While our approach explicitly considered NK cells, it can be generalized and is also applicable to other immunologically relevant populations such as T cells, etc. It is also clear from our approach that incorporating the dynamic response of both tumor and immune cells to further steps (e.g. from 50% to 25%) or longer phases (2 years) of dose reduction could further improve model predictions and ultimately lead to adaptive decisions based on individual responses.

A number of immune response-related CML models have been proposed, most of which use only tumor cell time courses to obtain optimal parameter fits and predict treatment response [33, 36, 38, 39]. Building on these models, we here provide new mechanistic insights, but also the concept that tracking the dynamic changes in a patient’s immunologic status over time can further improve model predictions. In a broader context, our model provides a flexible framework for studying tumor-immune interactions across different cancer types. Our findings not only advance the understanding of CML treatment dynamics, but also offer an important tool for quantitative, personalized medicine that integrates tumor-immune dynamics to optimize treatment decisions.

## Methods

### Model for tumor-immune interactions in CML treatment

We develop an ODE model for the interactions between tumor and immune effector cells during and after TKI treatment in CML patients (Fig. 1). The model considers active, TKI sensitive tumor cells (*T*), quiescent, TKI insensitive tumor cells (*T*_*N*_), and immune effector cells (*E*). Because there is biological evidence that quiescence is related to microenvironmental, niche-dependent effects, we denote quiescent tumor cells as niche-bound cells. By explicitly considering tumor-immune interactions at the cellular level, we generalize our previous work [37-39], and propose the following model

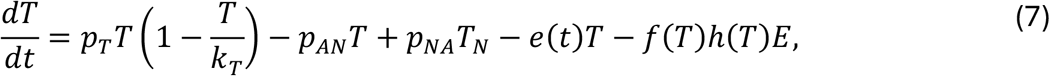

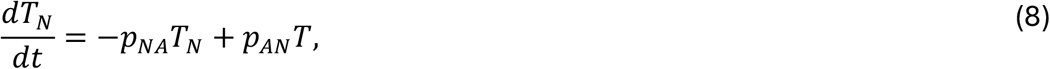

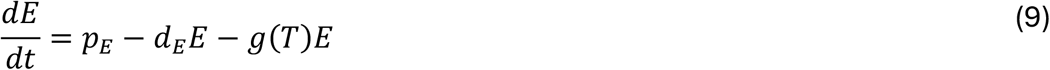

We assume that active tumor cells follow a logistic growth with net proliferation rate *p*_*T*_ and carrying capacity *k*_*T*_. They bind to the niche with rate *p*_*AN*_, and are eliminated with a treatment-related kill rate *e*(*t*) = *e*_*T*_ during treatment, while *e*(*t*) = 0 off-treatment. The killing of tumor cells by immune effector cells is described by −*f*(*T*)*h*(*T*)*E*. Here, *f*(*T*) is the number of tumor cells killed by each immune effector cell per unit of time in a normal, non-suppressed environment, while *h*(*T*) ∈ [0,1] accounts for a reduction in the number of functional immune cells due to the tumor-induced suppression of immune function. Both *f* and *h* depend on *T*, reflecting the modulation of immune activity and function by the tumor cells. While the immune activity *f*(*T*) is derived considering mechanisms of searching, targeting, and recharging (equation (1)), the immune function *h*(*T*) is derived from NK cell data (equation (2)). We further assume that quiescent, niche-bound tumor cells unbind with rate *p*_*NA*_, are not targeted by immune cells and insensitive to the treatment. Immune effector cells are produced at a constant rate *p*_*E*_ and die with rate *d*_*E*_, reaching a “healthy” steady state *p*_*E*_/*d*_*E*_ in absence of tumor cells. The suppression of immune cell recruitment by tumor cells is described by *g*(*T*), reflecting a tumor-dependent inhibition of the immune response. An expression for *g*(*T*) is derived using time courses of NK cell numbers (equation (3)). The model also describes tumor-immune interactions without a quiescence-inducing niche by neglecting equation (8) and setting *p*_*AN*_ = *p*_*AN*_ = 0. Initial numbers of immune and quiescent tumor cells are derived from quasi-steady state, *T*_*N*_(0) = (*p*_*AN*_/*p*_*NA*_)*T*(0) and *E*(0) = *p*_*Z*_/(*d*_*Z*_ + *g*(*T*(0)), while the initial number of active tumor cells *T*(0) is estimated from individual time courses.

### Mechanistic modeling of immune eflector cell activity

To model the immune cell activity against tumor cells, *f*(*T*), we revisit Holling’s work on predator-prey interactions [54, 55], associating tumor cells with prey and immune cells with predators. Holling assumed that the time *t*_*K*_ for a predator to kill a prey consists of a search time *t*_*S*_, and a handling time *t*_*K*_, during which the predator is occupied with its “meal”, including digestion (Fig. 1B). Holling assumed that the search time is inversely proportional to the prey density, since more prey means a faster search, while the handling time is constant, assumed as an intrinsic property of predator behavior.

Adapting to our context, the time it takes an immune cell to kill a tumor cell is *t*_*K*_ = *t*_*S*_ + *t*_*H*_ where *t*_*H*_ is constant and *t*_*S*_ = 1/(*sT*) where *s* is a proportionality constant related to the search speed; larger *s* results in faster searches. Therefore, the number of tumor cells killed by each immune effector cell per unit of time is

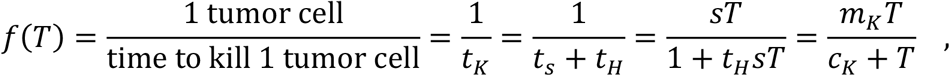

which is the classical Holling type-2 functional response (also a Michaelis-Menten kinetics) with saturation level *m*_*K*_ = 1/*t*_*K*_ and half-saturation constant *c*_*K*_ = 1/(*t*_*K*_*s*).

This functional response can also describe a more complex scenario considering a recovery time. Specifically, we assume now that immune cells target tumor cells by making “incursions” consisting of sequentially finding and eliminating several tumor cells, and then spending a fixed time to reactivate from an exhausted state (Fig. 1C). Thus, the incursion time consists of the sum of searching and handling times for each tumor cell, 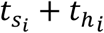, plus a final recharge time *t*_*R*_. Assuming a constant number of *m* tumor cells targeted in each incursion, we have

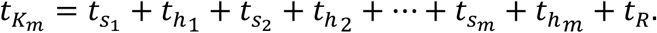

With searching times again inversely proportional to the number of tumor cells, we have

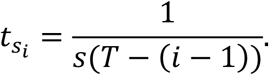

Assuming that the number of tumor cells targeted in each incursion is small compared to the total number of tumor cells (*T*− (*i* − 1) ≈ *T*), the individual searching times do not vary substantially and 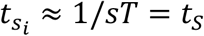. Further assuming equal handling times, we obtain

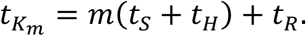

Thus, the number of tumor cells killed by each immune cell per unit of time is,

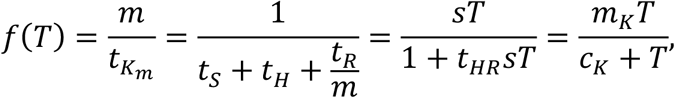

which is equivalent to the previous description, with the effective handling time *t*_*KR*_ = (*t*_*K*_ + *t*_*R*_/*m*) including a shared recharge time. This shows that a Holling type-2 functional response is suitable in the context of immune cells incursions and recharging time.

### Parameter estimation strategy

To reduce the number of patient-specific parameters and maximize model identifiability, we perform a dimensional analysis of system (7-9). We identify the non-dimensional groupings and make a sensible choice of which parameters can be fixed at the population level. Defining *x* = *T*_*N*_/*k*_*T*_, *y* = *T*/*k*_*T*_, *z* = *d*_*E*_*E*/*p*_*E*_ and *r* = *d*_*E*_*t*, system (7-9) becomes

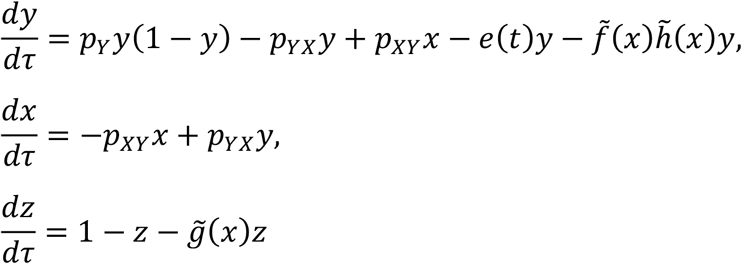

Where 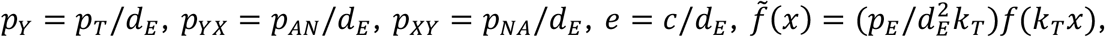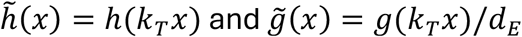

The death rate of immune cells *d*_*E*_ can be fixed if the growth and transition rates of tumor cells (*p*_*T*_, *p*_*NA*_ and *p*_*AN*_) and the production rate of effector cells (*p*_*E*_) can vary. Since the carrying capacity of tumor cells only scales the system size but does not change the model dynamics, it can also be fixed. Thus, without loss of generality, we assume population uniform values of *d*_*E*_ = 4 month^-1^ (mean lifetime of 1 week) and *k*_*T*_ = 10^6^ cells (reasonable for hematopoietic stem cells [56]).

Analyzing the model scale separations, we devised a stepwise approach that assigns the patient-specific parameters to different modules and estimates them as follows. First, we note that the parameters of *h*(*T*) (equation (2)) can be estimated from time course of NK cell functionality as a function of tumor load (Fig. 2A). Second, we observe that the dynamics of immune cells occurs on a fast timescale (Fig. S1A). Thus, the solution *E*(*t*) of (9) reaches the quasi-steady state *dE*/*dt* = 0 and is restricted to the slow manifold

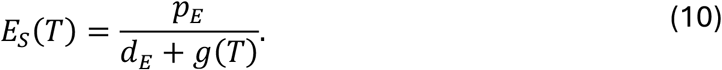

This refers to the set in the state space where the system dynamics is slower, and the number of immune cells is a function of the tumor load. In this case, equations (7) and (9) are replaced by a single equation implicitly considering immune effector cells,

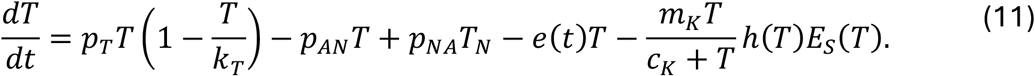

The slow manifold provides a good approximation for the solution *E*(*t*) of (9) (Fig. S1A) and was used to derive an expression for *g*(*T*) (equation (3)) and estimate its parameters and *p*_*E*_ from time courses of NK cell number as a function of tumor load (Fig. 2B, S1).

At this point, seven parameters remain, (*T*_0_, *p*_*NA*_, *p*_*AN*_, *e*_*TKI*_, *p*_*T*_, *m*_*K*_,*c*_*K*_) and are estimated as follows. The analysis of the effective immune response rate (equation (6)) showed that, due to the tumor-induced suppression, the effective immune-mediated kill of tumor cells is negligible during the initial treatment dynamics, independent of the values of *m*_*K*_ and *c*_*K*_ (Fig. 2D,3A). Therefore, in this step, the model can be approximated by the model without the immune system (equations (7-8) with *E* = 0). The resulting model is equivalent to our previous model [30], and its solution reproduces the typical biphasic decline described by a bi-exponential function, *T*_*biexp*_(*t*) = *Ae*^*αt*^ + *Be*^*βt*^. This simplified model can be solved analytically, allowing to derive an one-to-one correspondence expressing its parameters *p*_*NA*_, *p*_*AN*_, *q* = *e*_*T*_ − *p*_*T*_, *T*_0_ in terms of *A, B, α* and *β* [30]. Thus, for each patient time course with points (*t*_*i*_, *T*_*i*_), we fit the biexponential function *T*_*biexp*_(*t*) and use such correspondence to estimate parameters *p*_*NA*_, *p*_*AN*_, *q* = *e*_*T*_ − *p*_*T*_, *L*_0_. As shown in [30], this allows to estimate the net effect *q* = *e*_*T*_ − *p*_*T*_ resulting from treatment-induced cell death and tumor growth but does not disentangle such mechanisms; we choose to formulate the treatment effect as a function of the estimated value of *q* and the unknown value of *p*_*T*_, ie, *e*_*T*_ = *q*+ *p*_*T*_.

The remaining parameters to be estimated are *p*_*T*_, *m*_*K*_, *c*_*K*_, namely the tumor growth rate, the maximum immune kill rate, and the half-saturation constant of immune cell activity, respectively. Although they are critical for determining the outcome of treatment cessation, different combinations of *p*_*T*_, *m*_*K*_, *c*_*K*_ may yield equivalent dynamics under treatment with distinct outcomes after cessation (Fig. 3C,D). To integrate this variability in the choice of optimal parameters, we did not select a unique best fit but considered all parameter combinations that satisfy a given criterion for goodness of fit. Technically, we define a search set for each parameter based on reasonable ranges, *p*_*T*_ ∈ [0,1], *m*_*K*_ ∈ [0,1] and *log*_10_ *c*_*K*_ ∈ [1,4], the latter providing a range from MR5 to MR2 for the immune response threshold (Fig. 2D). Then, a Monte Carlo algorithm evaluates 10^6^ parameter combinations, and selects the 100 triples (*p*_*T*_, *m*_*K*_, *c*_*K*_) that minimizes the error between model and data [57, 58]. We found that *m*_*K*_ and *log*_10_ *c*_*K*_ were often correlated for each patient (Fig. S5AB). Therefore, we fixed *c*_*K*_ to a reasonable value of 10^2^ (Fig. 2D) and repeated the procedure for *p*_*T*_ and *m*_*K*_, now testing 10^4^ pairs, which resulted in equally good fits (Fig. S5C). Thus, to reduce parameter redundancy, the simplest approach of keeping *c*_*K*_ fixed was chosen. Applying this approach to tumor and NK cell time courses of the eight patients on full TKI dose from [13] resulted in excellent model fits, but with different predictions for treatment outcomes after discontinuation (Fig. 3C,D, S2). This illustrates that our stepwise approach is not only effective, but in a sense, it constrains all but two parameters with the information from clinical data, leaving the degrees of freedom only to the remaining parameters (*p*_*T*_, *m*_*K*_). It also shows that initially using the model without the immune system does not bias the estimates towards a weak immune system in the second step, as it also captures excellent fits predicting remission.

While the first cohort of eight patients with NK cell measurements contains time courses at full dose only, each of the 75 patient time courses from the DESTINY trial (see below) includes BCR-ABL1 measurements during an initial TKI treatment period at full dose, followed by a 1-year period at half dose, and a follow-up after treatment cessation. To take advantage of this data and assess how the predictive ability of the model increases with the inclusion of data under these different dose regimens, we applied the above approach in different settings increasing the selected data. First, the data consisted of measurements under full-dose only; second, we also included the half-dose period; third, we further included data points up to 6 months post-treatment discontinuation; and fourth, complete time courses were included. Since immune cell data were not available for these patients, we assumed population uniform values for the immune cell parameters *p*_*E*_ and *g*(*T*), using the mean values obtained from the eight data sets on NK cell response [13]. For each of the 75 patients, we obtained the 100 best fits as outlined above and defined the predicted probability of relapse at 12, 24 and 36 months after cessation as the fraction of fits predicting tumor load above MR3 (Fig. 4). We then compared the predictions to the reference true scenario, defined as the outcome predicted by the best fit using all data points. This was chosen instead of the data because it essentially described the data but also the extrapolated predictions.

## Data

Cytolytic NK cells (CD3^-^CD56^dim^CD1C^br^, see [13]) are a relevant population of immune effector cells and correlate with disease recurrence after treatment cessation. We utilize this population as a representative of immune effector cells described in our model. We use a dataset from [13], comprising time courses of eight CML patients undergoing standard TKI therapy, with measurements of NK cell number and function and tumor load at the same time points (Fig. 2A,B, S1, S2). The data on NK cell number consists of absolute numbers of CD3^-^CD56^dim^CD1C^br^ cells per microliter of peripheral blood, at diagnosis, pre-MMR, MMR, MR4.5, and TFR, while NK cell function is expressed as the percentage of CD107a^+^ NK cells at the same time points, normalized to control (healthy donors). The eight time courses of NK cell numbers were available from [13], while only the median values of the eight patients could be retrieved for the % of CD107a+ NK cells. Thus, the parameters regarding NK cell number (*p*_*E*_, *g*_*MAX*_, *c*_*R*_) were estimated from individual time courses, and the two parameters regarding NK cell function (*f*_*MIN*_, *c*_*F*_) are estimated from the median time course. The tumor load data is given as BCR-ABL1/ABL1 ratios (measured in %), which we assume to be proportional to tumor load, i.e., equal to 100 × (*T*/*k*_*T*_) [30]. Since this conversion implies that one tumor cell is equivalent to a BCR-ABL1/ABL1 ratio of 10^−4^%, undetectable tumors are set to 10^−5^%.

The second dataset includes the BCR-ABL1/ABL1 time courses of 75 of the 174 patients from the DESTINY trial [41], as previously used in [39]. See Figure 1A in that reference for details on patient selection criteria, relating to the exclusion of patients with no TKI reduction, recurrence during treatment, or undetectable tumor load measurements within the initial treatment period or initial measurements below MR3.

## Supplementary Information

Supplementary PDF file contains Figs S1-S5.

## Author contributions

Conceptualization: ACF, IR, IG. Formal Analysis, investigation, methodology, software, visualization, data curation: ACF. Writing – Original Draft Preparation: ACF, IG. Writing – Review & Editing: IR. Resources: AY, REC. Supervision: IG.

## Acknowledgments

ACF was supported by Alexander von Humboldt Foundation and Coordenação de Aperfeiçoamento de Pessoal de Nível Superior - Brasil (CAPES) - Finance Code 001, and partially supported by FAPEMIG RED-00133-21.

