## Supplementary Figures for "Predicting treatment-free remission outcomes in Chronic Myeloid Leukemia patients using an integrated model of tumor-immune dynamics"

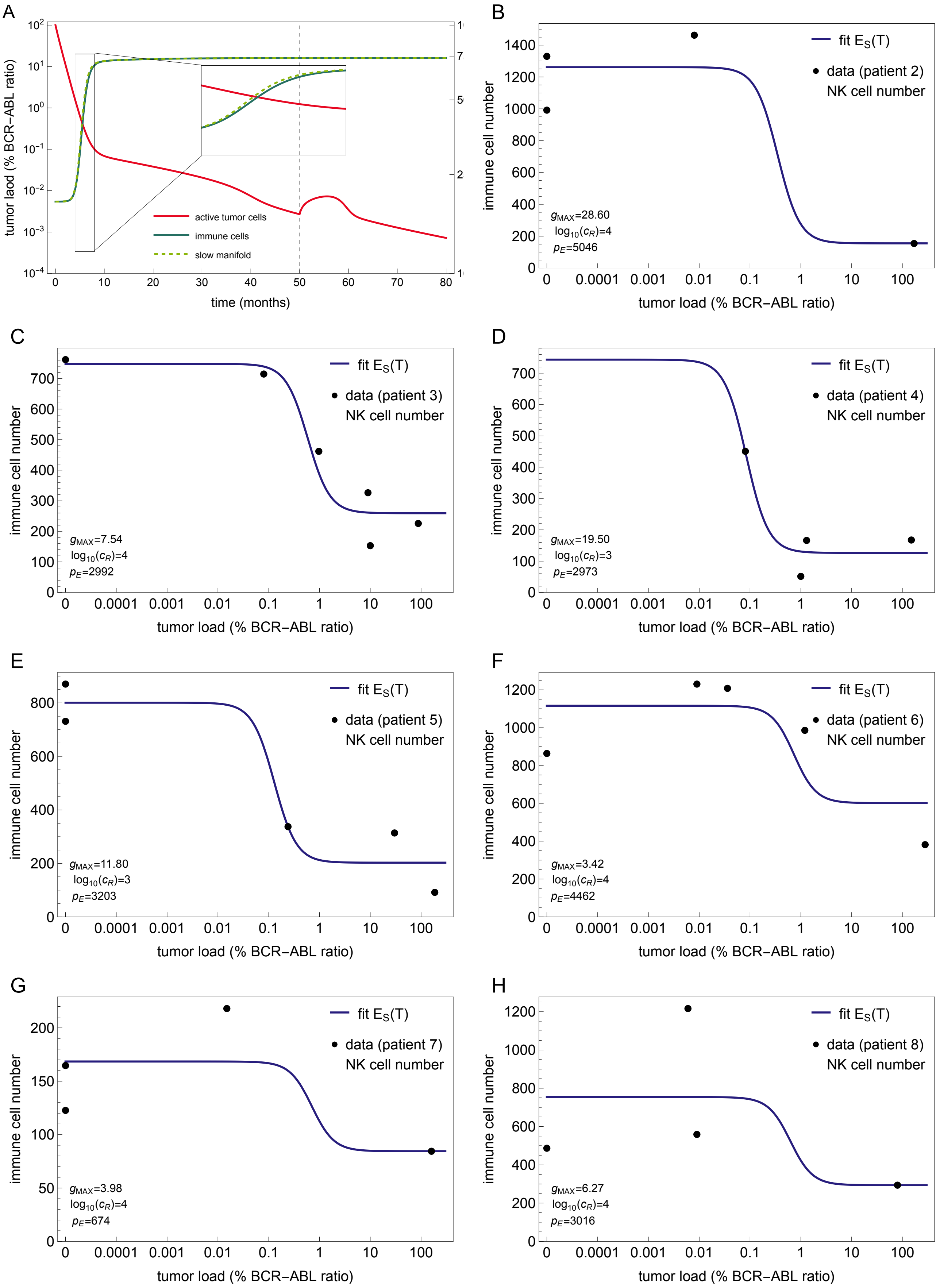


Supplementary Figure 1: A) Simulation with representative parameter values showing that the number of immune cells quickly reaches the quasi-steady state (equation (5), dashed green line). B-H) Fits of $E_S(T)$ (equation (9)) for other 7 patients with available NK cell time courses.

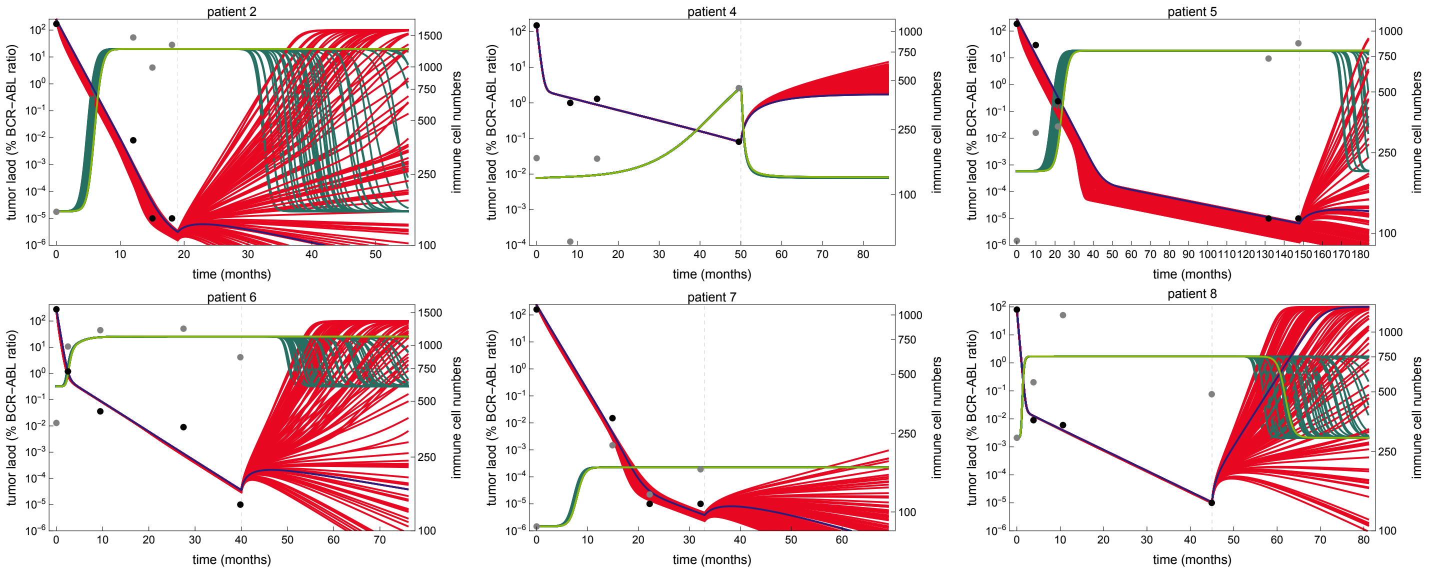


Supplementary Figure 2: Model fits. Model simulations of the 100 best fits to the time courses of tumor load (black dots) and NK cells (gray dots) for six representative patients. See also Figure 6.

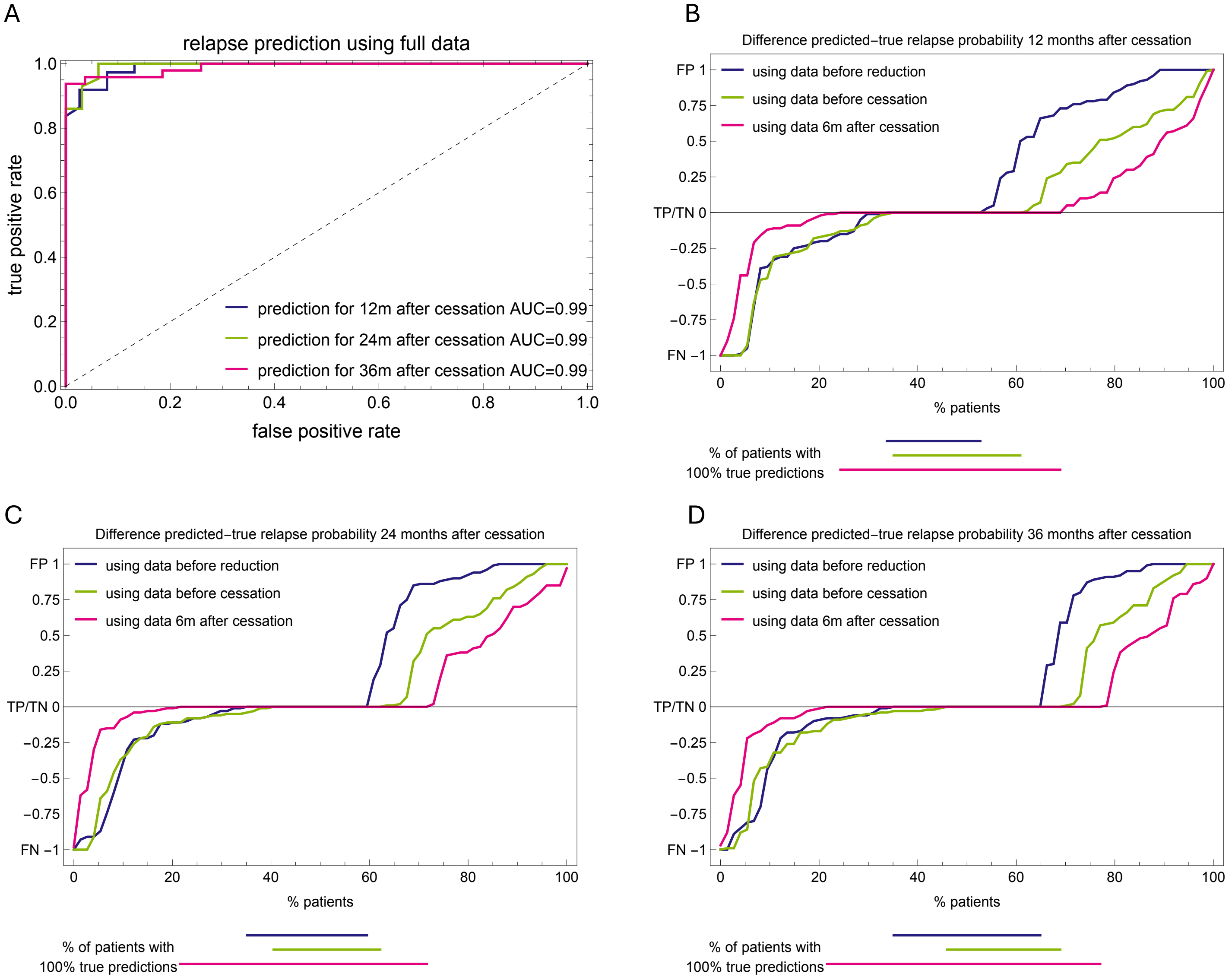


Supplementary Figure 3: A) Receiver operating characteristic (ROC) curves when the full time courses are used to fit the model. B,C,D) Differences between predicted and true recurrence probabilities using different data selections. Differences of 1, 0 and -1 correspond to false positives (FP), true positives/negatives (TP/TN) and false negatives (FN), respectively. The additional information contained in the data after dose reduction or cessation mainly changes the right part of these curves, corresponding to a correction of false positives.

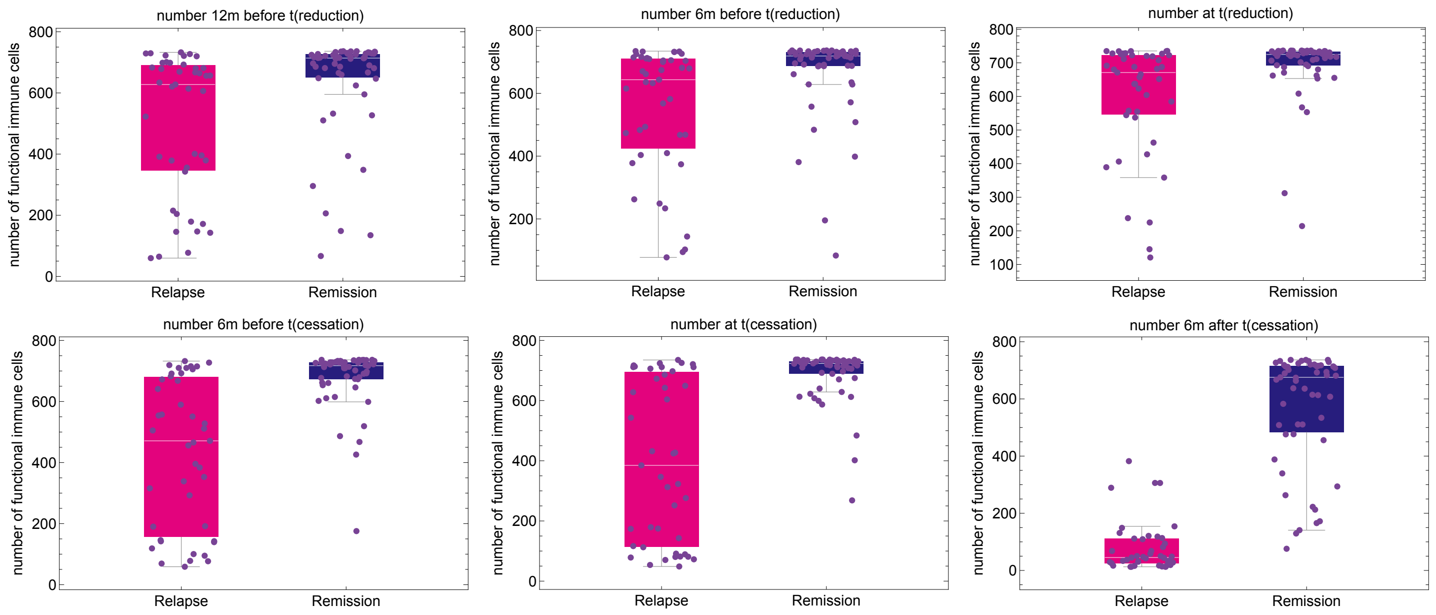


Supplementary Figure 4: The number of functional immune cells at different time points. A) Each plot shows the distribution of model estimates (mean values across all best fits) for the patient-specific number of functional immune cells at different time points before and after dose reduction or treatment cessation.

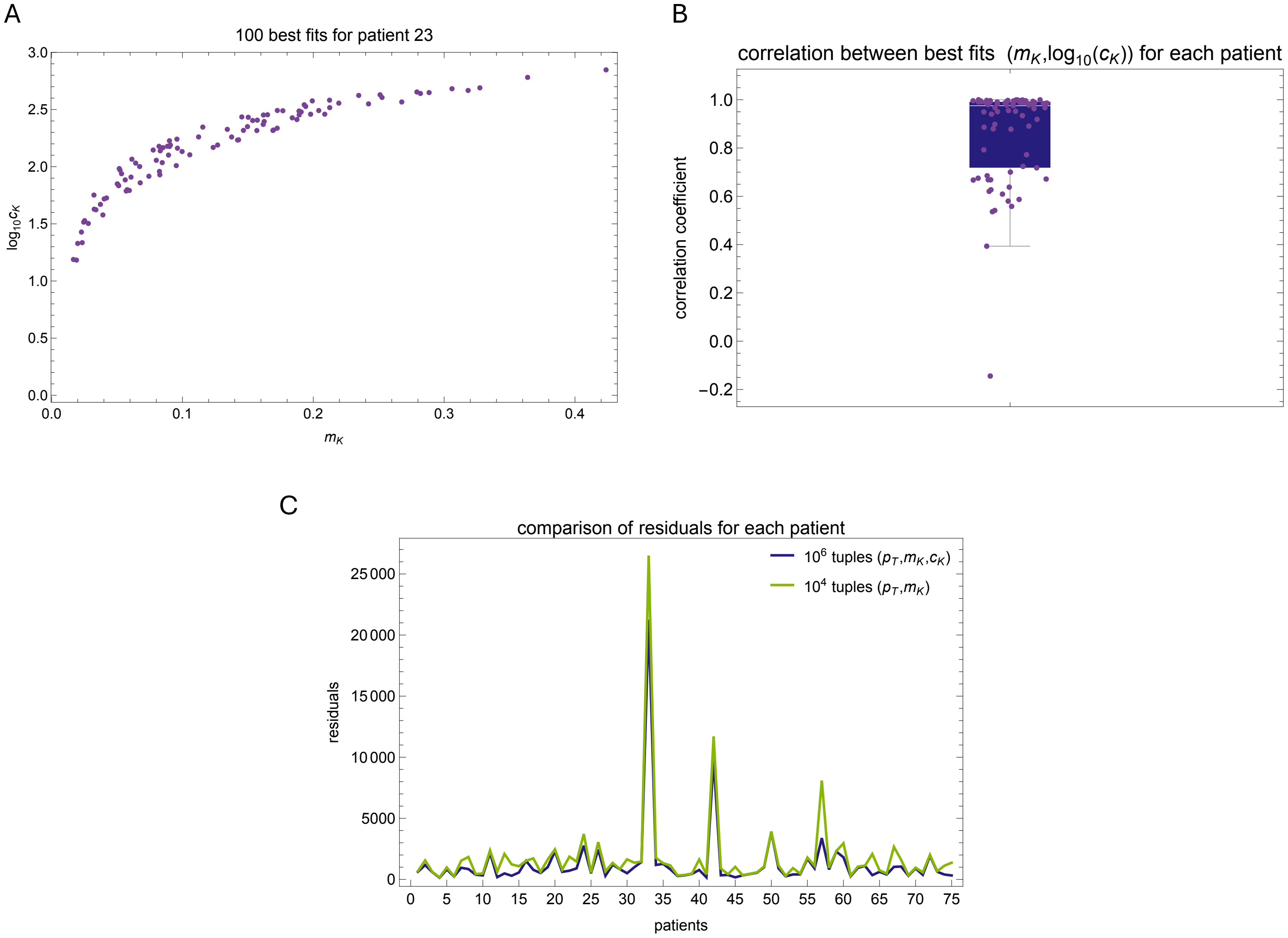


Supplementary Figure 5: A) Scatter plot showing the 100 best fits $(m_K, c_K)$ for patient 23 when the Monte Carlo procedure samples $10^6$ tuples of $(p_T, m_K, c_K)$; the Spearman correlation coefficient for this patient is 0.98. B) Distribution of the Spearman correlation coefficients for the 75 Destiny patients, with a median value of 0.96. C) Comparison of residuals using two procedures: sampling $10^6$ tuples of $(p_T, m_K, c_K)$ (purple) and sampling $10^4$ tuples of $(p_T, m_K)$ with $c_K = 10^2$ fixed (green). For each patient on the horizontal axis, the residuals are plotted on the vertical axis.
